# Rough and smooth variant *Mycobacterium abscessus* infections are differentially controlled by host immunity during chronic infection

**DOI:** 10.1101/856948

**Authors:** Elinor Hortle, Julia Y Kam, Elizabeth Krogman, Sherridan E Warner, Kaiming Luo, Tina Cheng, Pradeep Manuneedhi Cholan, Kazu Kikuchi, James A Triccas, Warwick J Britton, Matt D Johansen, Laurent Kremer, Stefan H Oehlers

## Abstract

Infections caused by *Mycobacterium abscessus* are increasing in prevalence within patient groups with respiratory comorbidities. Initial colonisation by the smooth colony *M. abscessus* (S) can be followed by an irreversible genetic switch into a highly inflammatory rough colony *M. abscessus* (R), often associated with a decline in pulmonary function. Our understanding of the role of adaptive immunity in *M. abscessus* pathogenesis is largely unknown. Here, we have used intraperitoneal infection of adult zebrafish to model *M. abscessus* pathogenesis in the context of fully functioning host immunity. We find infection with the R variant penetrates host organs causing an inflammatory immune response leading to necrotic granuloma formation within 2 weeks. The R bacilli are targeted by T cell-mediated immunity and burden is constrained. Strikingly, the S variant colonises host internal surfaces at high loads and is met with a robust innate immune response but little T cell-mediated immunity. Invasive granuloma formation is delayed in S variant infection compared to R variant infection upon which T cell-mediated immunity is required to control infection. In mixed infections, the S variant outcompetes the R variant. We also find the R variant activates host immunity to the detriment of S variant *M. abscessus* in mixed infections. These findings demonstrate the applicability of the adult zebrafish to model persistent *M. abscessus* infection and provide insight into the immunopathogenesis of chronic *M. abscessus* infection.

## Introduction

*Mycobacterium abscessus* is an increasingly recognized human pathogen responsible for a wide array of clinical manifestations including muco-cutaneous infections, disseminated or chronic pulmonary diseases (1). The latter is mostly encountered in patients with underlying lung disorders, such as bronchiectasis or cystic fibrosis (CF). Irrespective of being a rapid-growing mycobacteria (RGM), *M. abscessus* displays many pathophysiological traits with slow-growing mycobacteria (SGM), such as *Mycobacterium tuberculosis*. These include the capacity to persist silently within granulomatous structures and to produce pulmonary caseous lesions (2, 3). In addition, *M. abscessus* is notorious for being one of the most-drug resistant mycobacterial species, characterized by a wide panel of acquired and innate drug resistance mechanisms against nearly all anti-tubercular drugs, as well as many different classes of antibiotics (1, 4). Consequently, this explains the complexity and duration of the treatments and the high level of therapeutic failure (5).

*M. abscessus* exists either as smooth (S) or a rough (R) colony morphotype variants associated with distinct clinical outcomes (6). Previous epidemiological studies have highlighted the association of the R variant, persisting for many years in the infected host, with a rapid decline in pulmonary functions (7–9). It is well established that these morphological differences between S and R variants are dependent on the presence or absence of surface-exposed glycopeptidolipids (GPL), respectively (6, 10–12). However, our knowledge of the pathophysiological characteristics and interactions between R or S variants with the host immune cells remains largely incomplete and is hampered by the lack of animal models that are permissive to persistent *M. abscessus* infection (13).

Intravenous injection or aerosol administration of *M. abscessus* in immunocompetent BALB/c mice fails to establish a persistent infection, typified by a rapid clearance of the bacilli from the liver, spleen and lungs within 4 weeks (14). Immunosuppression is required to produce a progressive high level of infection with *M. abscessus* in mice, as shown in nude, SCID (severe combined immunodeficiency), interferon-gamma (GKO) and granulocyte-macrophage colony-stimulating factor (GM-CSF) knock-out mice (15).

The contribution of B and T cells in the control of *M. abscessus* infection has been studied in C57BL/6 mice with Rag2^-/-^, Cd3e^-/-^ and μMT^-/-^ knockouts (16). These studies indicated that infection control was primarily T cell dependent in the spleen, and both B and T cell dependent in the liver. In addition, IFNg-receptor KO mice (ifngr1^-/-^) were significantly impaired in their control of *M. abscessus* both in the spleen and in the liver, with markedly different granulomas and more pronounced in TNF^-/-^ mice (16). This points to the central role of T cell immunity, IFNg and TNF for the control of *M. abscessus* in C57BL/6 mice, similarly to the control of *M. tuberculosis* infection.

In recent years, alternative non-mammalian models, such as Drosophila (17), Galleria larvae (18), and zebrafish embryos (13) have been developed to study the chronology and pathology of *M. abscessus* infection and for *in vivo* therapeutic assessment of drugs active against *M. abscessus*. In particular, zebrafish embryos have delivered important insights into the pathogenesis of *M. abscessus* and the participation of innate immunity in controlling infection (10, 19). The optical transparency of zebrafish embryos has been used to visualise the formation of large extracellular cords by the R form *in vivo*, representing a mechanism of immune subversion by preventing phagocytic destruction and highlighting the importance bacterial virulence factors such as the dehydratase MAB_4780 and the MmpL8_MAB_ lipid transporter (10, 20, 21). Other studies in zebrafish embryos have demonstrated the contribution of host TNF signalling and IL8-mediated neutrophil recruitment for protective granulomatous immunity against *M. abscessus* (19), and the link between dysfunctional CFTR and vulnerability to *M. abscessus* infection via the macrophage oxidative response (22).

Adult zebrafish models have been well-described for the study of mycobacterial pathogenesis by *Mycobacterium marinum*, used as a surrogate for the closely related *M. tuberculosis*, and the human pathogen *Mycobacterium leprae* (23–26). Encompassing a fully functional immune system, previous studies in adult zebrafish with pathogenic mycobacteria such as *M. marinum* have unravelled the interplay between innate and adaptive immunity in mycobacterial granuloma formation and function.

Herein, we addressed whether adult zebrafish may be a useful host to analyse and compare the chronology of infection with *M. abscessus* S and R variants and to study the contribution of the T cell-mediated immunity and granulomatous response in *M. abscessus* infection.

## Results

### Adult zebrafish can be chronically infected with *M. abscessus*

We attempted to infect adult zebrafish with approximately 10^5^ CFU per animal with the rough (R) and smooth (S) variants of the reference strain CIP104536^T^, scaled for the smaller size of zebrafish from 10^6^-10^7^ used in mouse intravenous infections (14–16). To determine if *M. abscessus* produces a persistent infection in adult zebrafish, we performed CFU recovery on animals across 28 days of infection (Figure 1A). Variation in the initial inoculum ranging from 10^4^-10^6^ did not appear to impact the course of infection burden with stable burden of the R variant within a 1-log window either side of the inoculation dose to 28 days post infection (dpi) and progressive growth of the S variant to approximate 1-log above the inoculation dose at 28 dpi in all three experiments.

**Figure 1:**
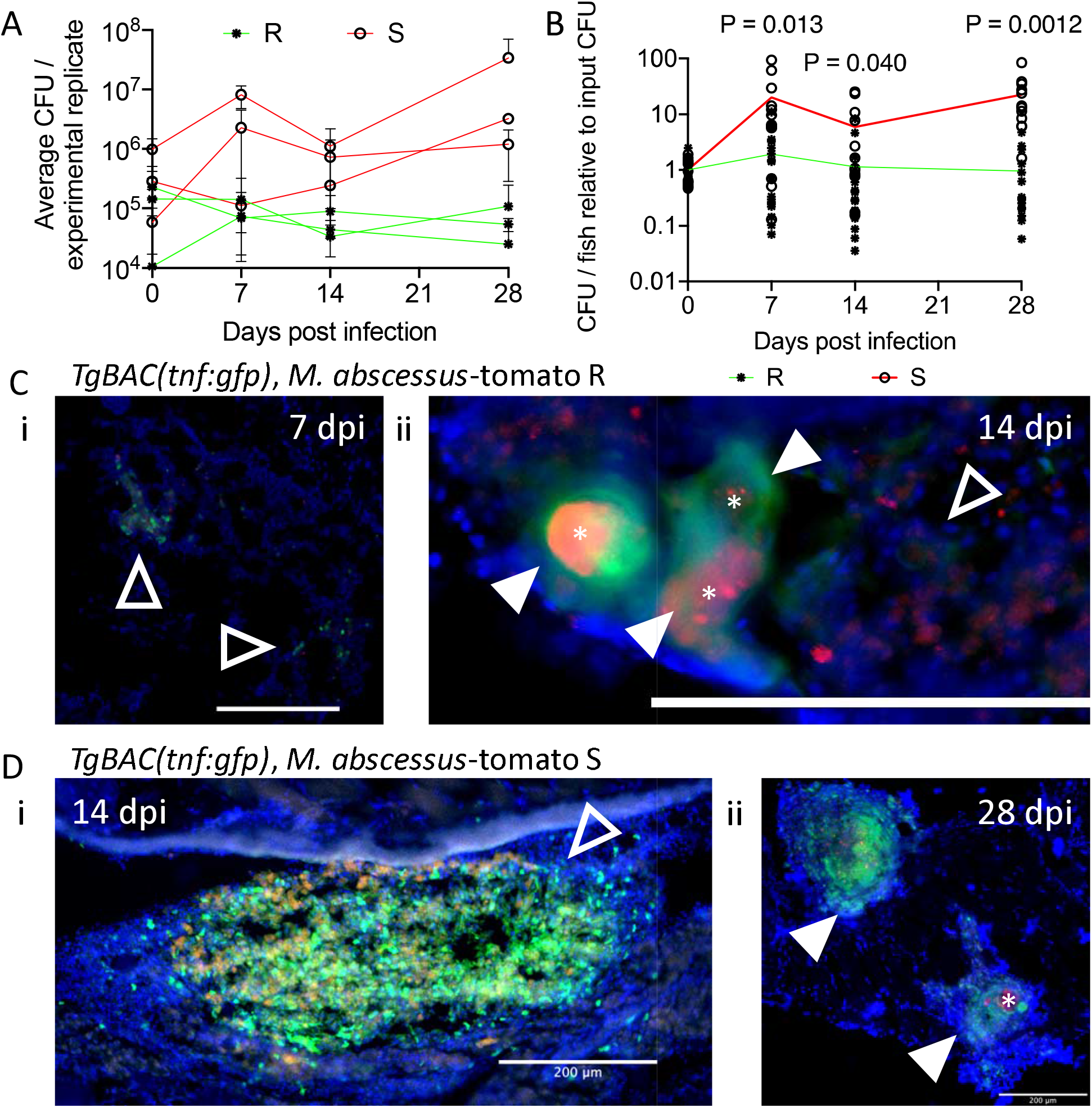
*M. abscessus* establishes chronic infection in adult zebrafish. A. Enumeration of CFUs from adult zebrafish infected with either the R or the S variant of *M. abscessus*. Each point represents a single experimental replicate with at least three animals per timepoint. Total n per timepoint: 0 dpi R=13 S=12; 7 dpi R=18 S=12; 14 dpi R=17 S=13; 28 dpi R=15 S=12. B. Relative CFUs recovered from adult zebrafish infected with either the R or the S variant of *M. abscessus*. Absolute CFU values were normalised to the inoculum CFU for each experimental replicate. Data is pooled from three replicates per *M. abscessus* variant. Total n per timepoint: 0 dpi R=13 S=12; 7 dpi R=18 S=12; 14 dpi R=17 S=13; 28 dpi R=15 S=12. Statistical tests by T test at each timepoint. C. Examples of R variant *M. abscessus*-tdTomato lesions in DAPI-stained cryosections from i. 7 dpi, and ii. 14 dpi *TgBAC(tnfa:GFP)^pd1028^* adult zebrafish. Filled arrowheads indicate organised necrotic granulomas, empty arrowheads indicate loose *M. abscessus* lesions, *tnfa* promoter induction is marked in green. D. Examples of S variant *M. abscessus*-tdTomato lesions in DAPI-stained cryosections from i. 14 dpi, and ii. 28 dpi *TgBAC(tnfa:GFP)^pd1028^* adult zebrafish. Filled arrowheads indicate organised granulomas, * indicate necrotic cores, empty arrowheads indicate loose *M. abscessus* lesions, *tnfa* promoter induction is marked in green. Scale bars indicate 200 μm.

Normalising burdens across three independent experiments per *M. abscessus* variant to perform statistical testing, we observed statistically significant increases in the proliferation of *M. abscessus* S compared to R at 7, 14, and 28 dpi (Figure 1B). Furthermore, comparison of the Day 0 and Day 28 burdens demonstrated *M. abscessus* R burdens were statistically unchanged (P>0.99, ANOVA) while *M. abscessus* S burdens increased by approximately 20x across the 4 weeks (P=0.024, ANOVA).

### Adult zebrafish mount a robust inflammatory response to *M. abscessus* infection

The cytokine *tumour necrosis factor* (*tnfa*) is essential for the granulomatous containment of *M. abscessus* in zebrafish embryos (19). To visualize *tnfa* transcription we next took advantage of the *TgBAC(tnfa:GFP)^pd1028^* zebrafish line (27), where GFP expression is driven by the *tnfa* promoter, to investigate if Tnfa expression is linked to granuloma formation. Expression of GFP was analysed in adult zebrafish infected with either variant of *M. abscessus* at 7, 14, and 28 dpi. GFP was expressed by host cells in close contact with *M. abscessus* both inside and outside of visibly organised granulomas (Figure 1C).

### Granuloma histopathology is accelerated during *M. abscessus* R infection compared to S

We next performed histology on adult zebrafish infected with fluorescent *M. abscessus* R or S. From 10 dpi we noted a heterogeneous mix of “unorganised lesions” visible as free bacteria around the peritoneal cavity or diffuse foci of bacteria spread throughout host tissue without concentric host nuclear organisation, and “organised granulomas” with stereotypical host nuclei ringing around a central focus of bacteria and the appearance of necrotic cores in all animals (Figure 2A).

**Figure 2:**
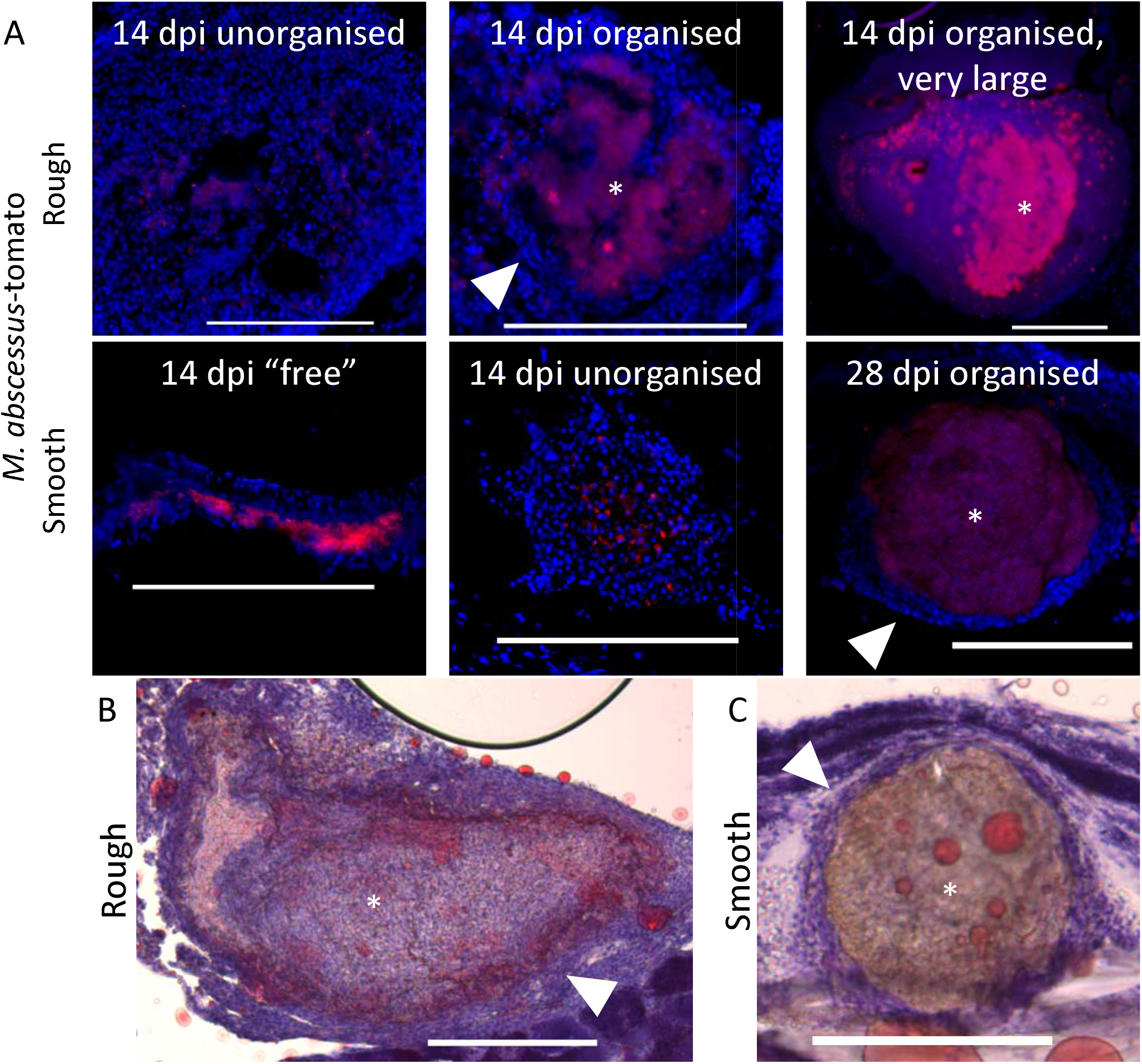
*M. abscessus* infection causes progressive granulomatous pathology. A. Stereotypical examples of bacterial lesions from DAPI-stained sections from adult zebrafish infected with *M. abscessus* expressing tdTomato. Top row infected with the rough variant, bottom row infected with the smooth variant, timepoint as indicated. B. Example of Oil Red O-stained very large granuloma in a 14 dpi adult zebrafish infected with R *M. abscessus*. Neutral lipid staining is indicated by red colouration in the macrophages layer surrounding the mycobacterial core, host nuclei are counterstained purple with haematoxylin. C. Example of Oil Red O-stained large granuloma in 28 dpi adult zebrafish infected with S *M. abscessus*. Note lack of lipid staining in the macrophage rim of the granuloma compared to R granuloma. Scale bars indicate 200 μm. Filled arrowheads indicate epithelised macrophage nuclei forming a stereotypical concentric layer surrounding the mycobacterial core of organised granulomas, * indicate necrotic cores.

We also observed the appearance of very large granulomas filled with fluorescent bacteria and necrotic debris measuring over 500 μm in *M. abscessus* R-infected animals from 14 dpi onwards (Figure 2A). These large granulomas were observed only occasionally and at a rate of no more than 1 per infected animal at 14 and 28 dpi (n=2 with single abscess, 9 without abscess). Three *M. abscessus* R-infected animals were maintained until 70 dpi, all three were found to multiple have granulomas containing fluorescent *M. abscessus* R demonstrating very long lasting infection is possible in adult zebrafish, and two were found to have granulomas measuring over 500 μm suggesting an increase in large granuloma formation with infection duration (Supplemental Figure 1). Granulomas in *M. abscessus* S-infected animals did not reach this size at the 14 and 28 dpi timepoints sampled (n=15 without abscess).

Oil red O staining revealed the accumulation of foam cells in cellular rim of *M. abscessus* R granulomas (Figure 2B), consistent with immunopathology seen in immunocompromised mice infected with *M. abscessus* (15). Conversely, there was little Oil Red O staining in *M. abscessus* S granulomas indicating a lack of foam cell formation (Figure 2C).

We next quantified the number of lesions with the categories of “organised granulomas”, with host nuclear organisation into rings, and “unorganised lesions”, consisting of either diffuse foci of bacteria spread throughout host tissue or free bacteria around the peritoneal cavity at the whole animal level. The proportion of “organised granulomas” in *M. abscessus* R-infected adult zebrafish from appeared to increase after 10 dpi, however this change was not statistically significant (Figure 3A). *M. abscessus* S was observed to grow freely in mesenteric spaces and form poorly organised cellular granulomas at 14 dpi before the proportion of “organised granulomas” increased at 28 dpi (Figure 3A). The proportion of “organised granulomas” was higher in *M. abscessus* R-infected than in *M. abscessus* S-infected animals at 14 dpi suggesting granuloma formation is accelerated in R infections compared to S.

**Figure 3:**
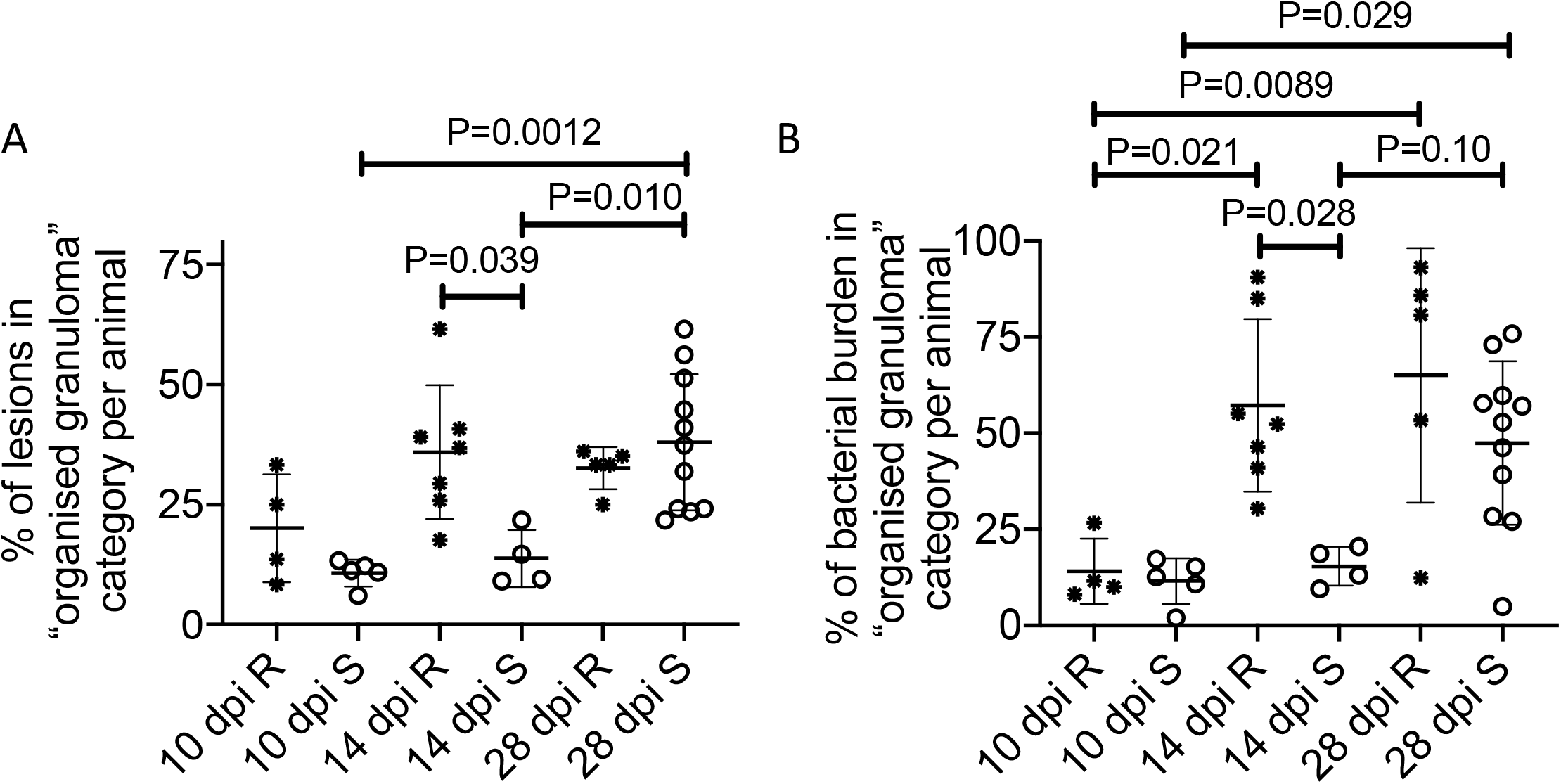
Granuloma histopathology is accelerated during *M. abscessus* R infection compared to S. A. Quantification of bacterial lesion organisation in adult zebrafish infected with approximately 10^5^ CFU *M. abscessus*. B. Quantification of bacterial burden stratified by lesion organisation in adult zebrafish infected with approximately 10^5^ CFU *M. abscessus*. Total individual lesions analysed (organised/unorganised): 10 dpi R (37/228); 10 dpi S (107/788);14 dpi R (180/314); 14 dpi S (61/352); 28 dpi R (49/93); 28 dpi S (316/476). Statistical testing by ANOVA.

These patterns were recapitulated in our quantification of fluorescent bacterial burden in each type of lesion. Significantly more *M. abscessus* R was observed within “organised granulomas” at 14 and 28 dpi than at 10 dpi, and an increase in the proportion of *M. abscessus* S within “organised granulomas” was only observed at 28 dpi compared to 10 and 14 dpi (Figure 3B). The proportion of *M. abscessus* R within “organised granulomas” was higher than the proportion of *M. abscessus* S within “organised granulomas” at 14 dpi, again suggesting accelerated granuloma formation in R variant infections compared to S.

### T cell-dependent immunity differentially controls infection by *M. abscessus* variants

Given the requirement for T cells to maintain granuloma structure in adult zebrafish *M. marinum* infection (25), we next asked if there was T cell involvement around *M. abscessus* granulomas using *TgBAC(lck:EGFP)^vcc4^* zebrafish (28). We observed T cell association and penetration throughout unorganised and organised *M. abscessus* R granulomas, but T cells were largely excluded from the cores of the very large abscess-like lesions (Figure 4A). We failed to observe T cell interaction with *M. abscessus* S growing “free” around peritoneal organs early in infection, furthermore the T cell response to tissue-invasive *M. abscessus* S was noticeably less than that for equivalent sized *M. abscessus* R granulomas (Figure 4B and Supplemental Figure 2).

**Figure 4:**
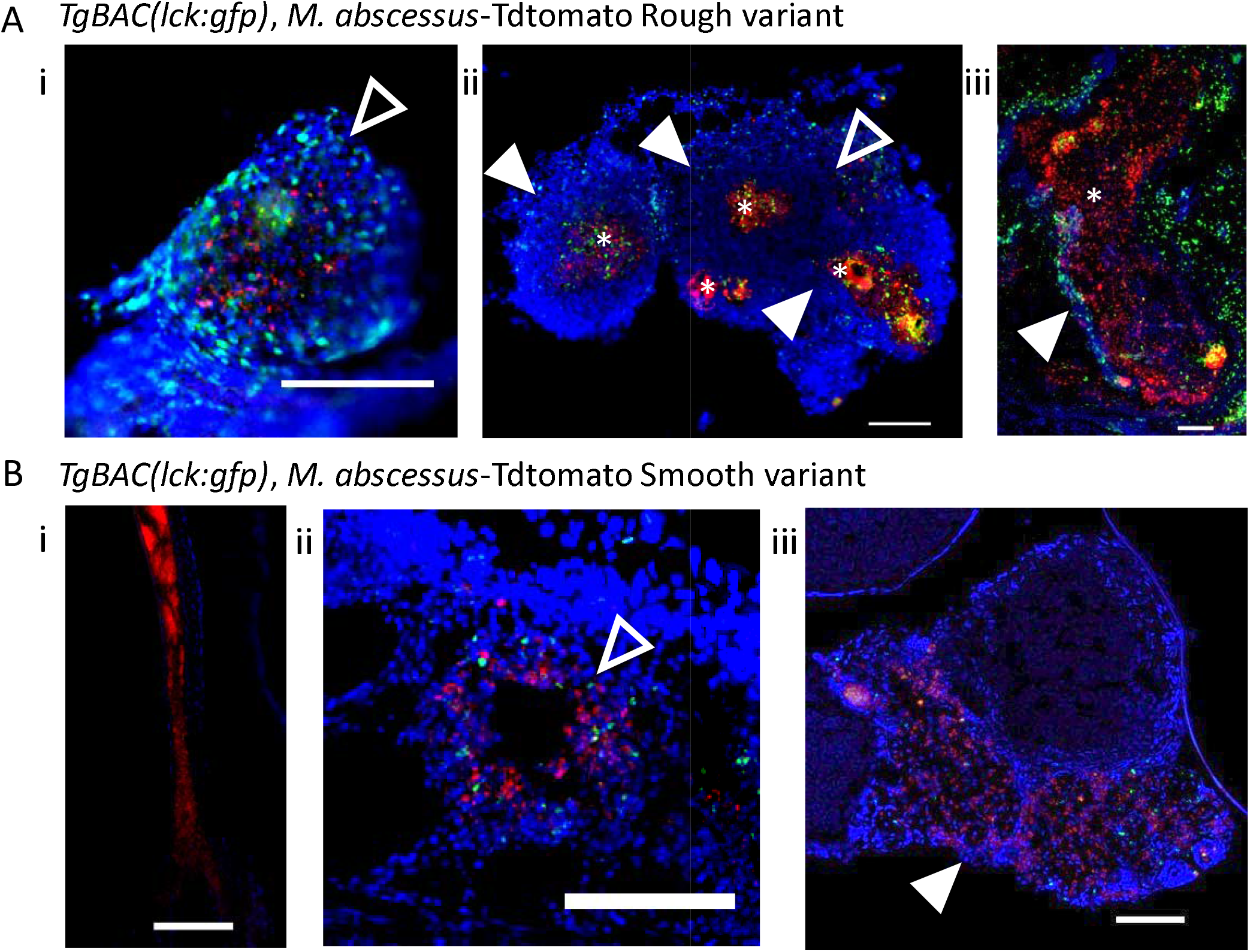
T cell recruitment to S *M. abscessus* infection is delayed compared to R. A. Examples of T cell recruitment to granulomas in 14 dpi *TgBAC*(*lck*:*EGFP*)^*vcc4*^ adult zebrafish infected with R *M. abscessus*-tdTomato. i. Example of an unorganised granuloma. ii. Example of a multilobed organised granuloma. iii. Example of a very large granuloma. B. Example of lack of T cell recruitment to S *M. abscessus*- tdTomato in 14 dpi *TgBAC(lck:EGFP)^vcc4^* adult zebrafish. i. Example of *M. abscessus* S mass growing “free” in peritoneal cavity. ii. Example of an unorganised granuloma. iii. Example of an organised granuloma. Scale bars indicate 100 μm. Filled arrowheads indicate organised granulomas, * indicate necrotic cores, empty arrowheads indicate loose *M. abscessus* lesions, *lck:gfp* positive T cells are marked in green.

To directly test the requirement of T cells in containing *M. abscessus* we next utilised the *lck^-/-sa410^* mutant line which is T cell-deficient. We infected wild type (WT) control and *lck^-/-sa410^* mutant adult zebrafish with both the S and R variants. T cell-deficient adult zebrafish were significantly more susceptible to *M. abscessus* R infection with reduced survival over 28 days of infection (P = 0.0005, Log-rank test) (Figure 5A). T cell deficiency had a less pronounced effect on the survival of animals infected with *M. abscessus* S compared to *M. abscessus* R infection (WT S versus lck-/- S P = 0.03, Log-rank test). Within the T cell-deficient animals, there was a 5.5 day increased median survival for *M. abscessus* S-infected animals (34 dpi) compared to *M. abscessus* R (28.5 dpi), although both groups eventually succumbed to infection at the same rate after 35 dpi (P = 0.78, Log-rank test).

**Figure 5:**
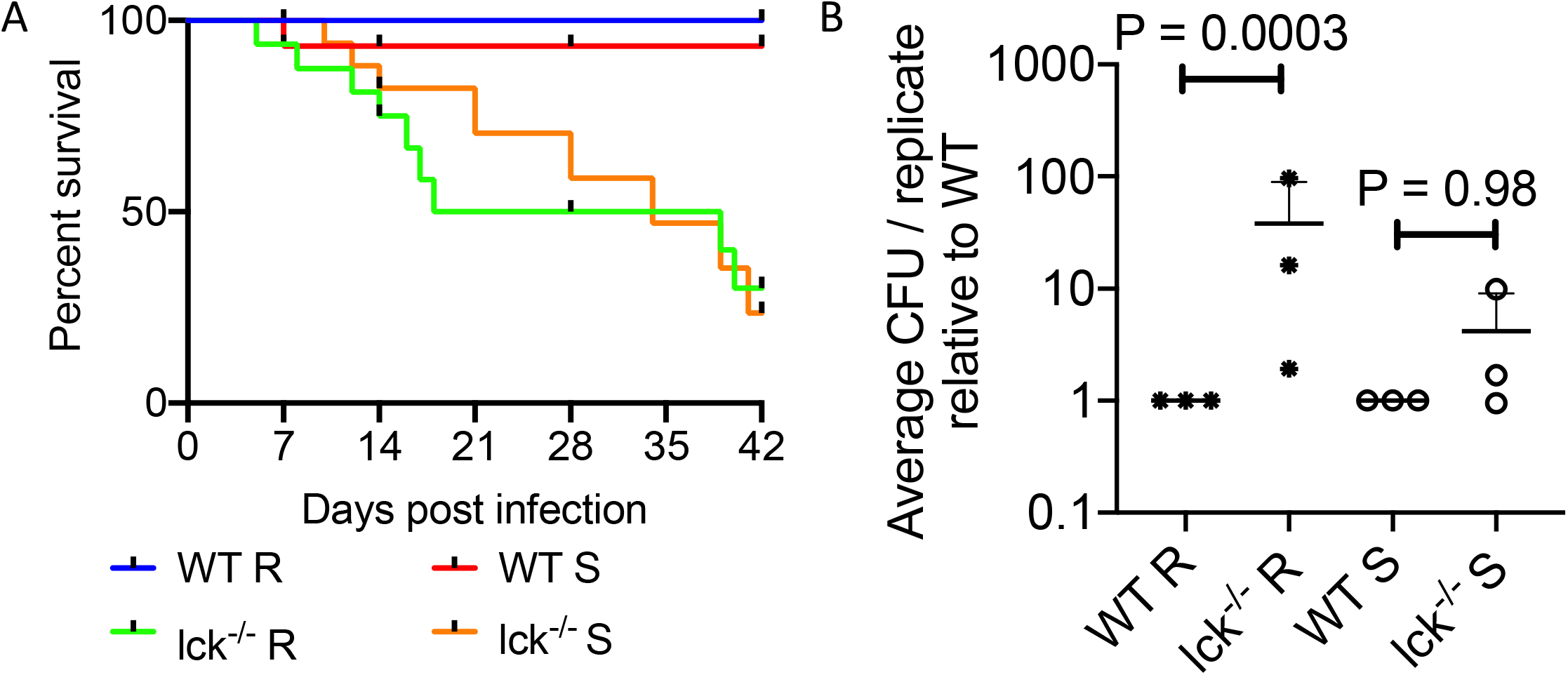
T cells are necessary to control R but not S *M. abscessus* infection. A. Survival analysis of WT and lck^-/-*sa410*^ adult zebrafish infected with R or S *M. abscessus*. Total n=12 *WT/Mabs* R; 16 lck-/-/*Mabs* R; 15 *WT/Mabs* S; 22 lck-/-/*Mabs* S. B. Normalised CFUs recovered from 14 dpi WT and lck^-/-*sa410*^ adult zebrafish infected with *M. abscessus*. Each point represents the average of a single experiment with at least 2 animals per group. Total n per group: R WT=12 lck^-/-^=15; S WT=11 lck^-/-^=12. Statistical testing by 2-way ANOVA.

Surprisingly, we found necrotic granulomas in a survivor 56 dpi *lck^-/-sa410^* fish infected with *M. abscessus* R (Supplemental Figure 3). These granulomas were all relatively small compared to the large granulomas seen in our small sample of 70 dpi WT animals with the largest having necrotic cores of approximately 100 μm.

Bacterial burden was significantly increased in 14 dpi *lck^-/-sa410^* animals infected with the R, but not the S variant compared to burdens in WT adult zebrafish (Figure 5B). These observations suggest the initial control of *M. abscessus* R infection is more reliant on T cell-mediated immunity than the control *M. abscessus* S infection during the first 2-3 weeks of infection.

### The *in vivo* survival advantage of *M. abscessus* S is compromised by *M. abscessus* R in mixed infections

To further examine our observation that *M. abscessus* S has a survival advantage over *M. abscessus* R in the adult zebrafish infection model, we performed co-infection of adult zebrafish with equal numbers of each variant expressing either Wasabi or tdTomato fluorescent proteins to enable simple tracking (Figure 6A). Coinfection did not affect the recovered *M. abscessus* R burden as near identical *M. abscessus* R CFUs were recovered from single and mixed-infected animals at 7 dpi (Figure 6B). However, coinfection did cause a decrease in the number of recoverable *M. abscessus* S from the levels found in single infections demonstrating a negative effect of R infection on the survival of *M. abscessus* S (Figure 6B). Despite this drop in S burden, analysis of the ratio of recovered R:S colonies revealed a clear and rapid shift in population proportions from 1:1 at 1 dpi to 0.5 rough:1 smooth ratio at 7 dpi that remained stable through to 14 dpi demonstrating partial retention of the relative survival advantage in mixed infection (Figure 6C).

**Figure 6:**
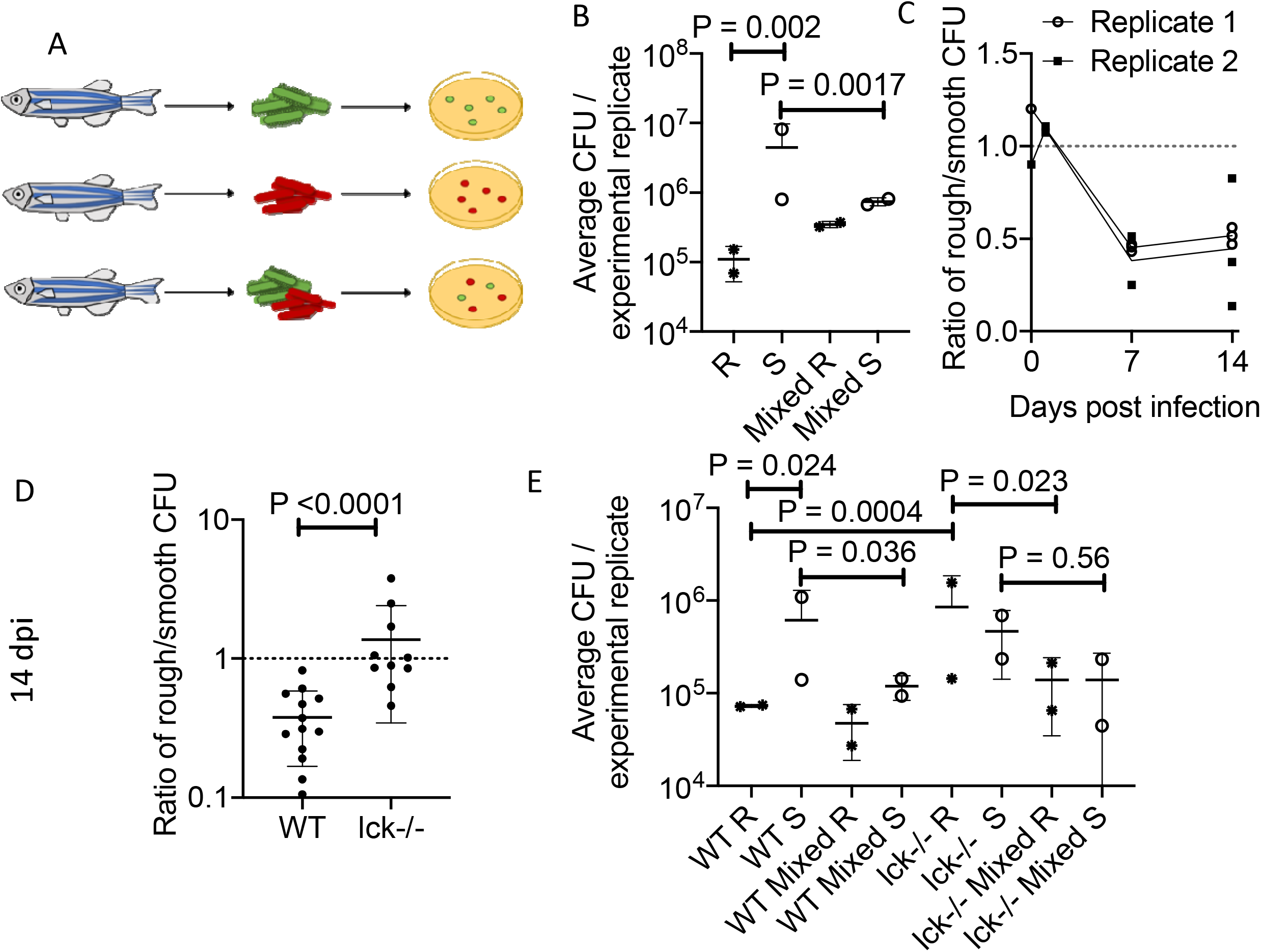
The *in vivo* survival advantage of *M. abscessus* S is compromised by *M. abscessus* R in mixed infections. A. Schema outlining mixed infection experiment. B. Enumeration of CFUs 7 dpi WT adult zebrafish outlined in panel A. Each point represents the average of a single experiment with at least 2 animals per group. Total n per group: Single R=8, Single S=5, Mixed=5. Statistical testing by 2-way ANOVA. C. Ratio of R:S CFUs recovered from individual WT adult zebrafish from the mixed infection group outlined in panel A. D. Ratio of R:S CFUs recovered from 14 dpi WT and lck^-/-*sa410*^ adult zebrafish infected with a mixture of differentially labelled R and S variants. Statistical testing by Mann-Whitney test. E. Enumeration of CFUs from WT and lck^-/-*sa410*^ adult zebrafish divided into the three groups outlined in panel A. Each point represents the average of a single experimental with at least 3 animals per group. Total n per group: WT R=12 S=10 Mixed=10, lck^-/-^ R=9, S=8, Mixed=10. Statistical testing by 2-way ANOVA.

We hypothesised that the T cell response induced by the R variant to antigens shared with the S variant could be responsible for the clearance of S variant in mixed infections. Using our mixed infection animals as individually-controlled experiments, we compared the ratio of R:S colonies at 14 dpi in WT and *lck^-/-sa410^* zebrafish (Figure 6D). We observed a higher R:S ratio in *lck^-/-sa410^* zebrafish than in WT zebrafish, suggesting a role for either 1) T cell-dependent immunity in controlling *M. abscessus* R growth or 2) enhanced T cell-independent immunity in suppressing *M. abscessus* S growth.

To distinguish between these two possibilities, we analysed the recovered CFU from WT and T cell-deficient *lck^-/-sa410^* animals with single and mixed infections of R and S *M. abscessus* at 14 dpi (Figure 6E). CFU recovery from these mixed infections in the T cell-deficient animals revealed a complex interaction between T cell depletion and *M. abscessus* burdens in mixed infections compared to single variant infections. The burden of the R variant *M. abscessus* recovered from mixed infections was significantly lower than the burdens achieved in single infections, suggesting a T cell-independent effect of the S variant on restraining R growth (P=0.023, 2 way ANOVA). However, there was only a non-statistically significant trend to increased *M. abscessus* R in *lck^-/-sa410^* mutants compared to WT animals (P=0.99, 2-way ANOVA). Finally, we observed a reduction in the S variant burden recovered from mixed infections compared to single infections in T cell-deficient animals (Figure 6E). However, there was no difference between the recovery of S variant *M. abscessus* in WT or *lck^-/-sa410^* mutants (P<0.99, 2 way ANOVA), highlighting the complex role of T cells in the R-triggered clearance of S variant *M. abscessus*.

## Discussion

In this study, we report for the use of adult zebrafish to probe both host and mycobacterial determinants of pathogenesis during persistent infection with *M. abscessus*. Infection with the R and S variants was maintained at high levels up to one month post infection in genetically intact animals, a major improvement on traditional mouse models of *M. abscessus* infection.

While the R variant induces a more robust and aggressive infection than the S variant in zebrafish embryos (10), this appears to not be the case in the adult fish. We observed better control of R variant burden and establishment of a higher burden of persistent infection with the S variant. One possible explanation for the better survival of the S compared is that infection with the R variant results in earlier granuloma formation and engagement of T cells. This contribution of the T cell response was further substantiated using T cell-deficient fish, where infection of *lck* fish with the R bacilli resulted in a higher bacterial burden than in WT fish at 14 dpi, which was not observed with S bacilli. These observations provide insight into the clinical observation that AIDS patients are not at increased risk of *M. abscessus* infection to the same degree that AIDS is a risk factor for *M. tuberculosis* and other non-tuberculous mycobacterium infections such as *Mycobacterium avium* (29).

It is well known that the intracellular lifestyle of the R and S morphotypes differ significantly, resulting in entirely distinct infection scenarios that, we hypothesise, underlie the accelerated granuloma formation by the R variant in adult zebrafish (30). The absence of GPL on the outer mycomembrane causes corded growth of R variants, resulting in multiple bacilli being simultaneously phagocytosed by macrophages and overloaded phagosomes that rapidly activate autophagy pathways (12, 30). Comparatively, the S variant is able to survive for an extended period of time within the phagosome, producing a chronic and persistent infection (31). As such, these polar infection responses may explain why the R variant displays widespread organised granuloma formation by 14 dpi, compared to S which shows a delayed onset of granuloma formation after 14 dpi. Moreover, this observation matches the superior *in vivo* growth performance of S bacilli compared to R, suggesting that the R variant is at an overall disadvantage because of its intrinsic hyper-inflammatory status and the activation of T cell-mediated immunity that appears concomitant with granuloma formation. Interestingly, earlier reports using the zebrafish embryo demonstrated that both bacterial burden and granuloma formation dynamics were similar between both the S and R variants (10, 22), highlighting the critical role of adaptive immunity in divergent *M. abscessus* infection responses. Taken together, our data provide additional evidence for the distinct intracellular fates of both S and R variants *in vivo*, and further implicates the role of adaptive immunity in granuloma formation and control of *M. abscessus* infection in an adult zebrafish model.

T cells are critical host determinants in the control of mycobacterial infection (29). Recruitment of T cells into granulomas is thought to be essential in containing persistent infection, while T cell deficiencies are associated with greater mycobacterial infection severities (23, 29, 32, 33). Recently, an adult zebrafish infection model for *M. leprae* demonstrated that T cells are essential for containment of infection (23). Herein, we examined the recruitment of T cells within granulomas and identified that S variant granuloma were marked by a relative paucity of T cell infiltration, suggesting that T cells may play a less significant role in S variant infections than those with R variants. Using the *lck^-/-^* zebrafish, we observed displayed an improved *in vivo* growth performance of the R variant in the absence of T cells when compared to WT animals, highlighting the role of T cells in the control of R variants. This observation was not maintained with the S variant, which showed no increase in bacterial growth *in vivo* irrespective of the absence of T cells early in infection, despite *lck*-/- fish succumbing to intraperitoneal infection within 40 days at the same rate in the absence of T cells irrespective of bacterial morphotype.

Our co-infection experiments further support the theory that tissue destruction caused by the R variant activates protective trans-acting host immunity that impairs further *M. abscessus* growth. This was seen most clearly in the restriction of *M. abscessus* S growth in mixed infections. It suggests *M. abscessus* must balance the benefits of R variant pathogenicity allowing individuals to kill and escape macrophage containment, with the need to avoid activation of host-protective immunity at a population level when adapting to an animal host. Although we clearly observed an equalisation of this effect in T cell-deficient mutants (Figure 6D), we were unable to determine if this was due to an increase in R growth or suppression of S growth (Figure 6E).

The extended maintenance of R variant burden for at least 4 weeks in zebrafish is comparable to our recent data from the C3HeB/FeJ “Kramnik” mouse (34), but the proliferation of S variant up to a log above inoculation dose is unprecedented in a genetically intact vertebrate host. The granulomatous immunopathology in mycobacterium-infected C3HeB/FeJ mice is due to an exaggerated type I interferon response suppressing protective interleukin-1 (35). Further analysis of interferon and interleukin-1 responses to *M. abscessus* infection of mice and zebrafish will help translate our understanding of these dichotomous responses into host directed therapies.

We did not observe switching of S *M. abscessus* into a rough colony morphotype at any timepoint during this or subsequent studies. *In vivo* switching is a rare event that has only been documented in immunocompromised mice or after months-to-years in patients (36, 37). The high S morphotype burdens achieved in adult zebrafish suggest this platform may be useful for future studies of switching during extended infections, with the potential to model responses to chemotherapy.

To date, our understanding of the diverse immune responses between S and R variants have essentially been thoroughly described with respect to innate immunity, and currently our knowledge pertaining to adaptive immunity in *M. abscessus* infection has been poorly characterised (16). Using this new adult zebrafish *M. abscessus* infection model, we have shown that S and R variants produce strikingly different disease phenotypes, which were further exemplified in the absence of T cells. Consequently, these results suggest that the host-pathogen interactions dictating *M. abscessus* pathogenesis are complex and implicate adaptive immunity to a greater extent than originally anticipated. Future work should exploit this relevant animal model in combination with zebrafish lacking the cystic fibrosis transmembrane conductance regulator (CFTR) gene, and for the development and testing of novel antibiotics and vaccine candidates that may be used for the treatment of *M. abscessus* infection.

## Methods

### Zebrafish strains and handling

Zebrafish strains used in this study are AB strain wildtype, *TgBAC(tnfa:GFP)^pd1028^, TgBAC*(*lck*:*EGFP*)^*vcc4*^, *lck-/-^sa410^* (27, 28) between 3 and 6 months of age. Animals were held in a 28°C incubator with a 14:10 hour light:dark cycle. Animals were infected by intraperitoneal injection with approximately 10^5^ CFU *M. abscessus*, unless otherwise stated, using a 31 G insulin needle and syringe as previously described (38). Infected zebrafish were recovered into system water and held in 1 L beakers with daily feeding for the duration of the experiment. Infection experiments were carried out with ethical approval from the Sydney Local Health District Animal Welfare Committee approval 16-037.

### *M. abscessus* strains and handling

Rough (R) and smooth (S) variants of *M. abscessus* strain CIP104536^T^ were grown at 37°C in Middlebrook 7H9 broth supplemented with 10% Oleic acid/Albumin/Dextrose/Catalase (OADC) enrichment and 0.05% Tween 80 or on Middlebrook 7H10 agar containing 10% OADC (7H10 OADC). Recombinant *M. abscessus* strains expressing tdTomato or Wasabi were grown in the presence of 500 μg/ml hygromycin (10, 19). Homogenous bacterial suspensions for intraperitoneal injection in adult fish were prepared as previously reported (39).

### Bacterial recovery

Animals were euthanised by tricaine anaesthetic overdose and rinsed in sterile water. Individual carcasses were homogenised and serially diluted into sterile water. Homogenates were plated onto 7H10 supplemented with OADC and 300 μg/ml hygromycin. Plates were grown for at least 4 days at 37°C.

### Histology

Animals subjected to cryosectioning as previously described (38). Briefly, euthanasia was performed by tricaine anaesthetic overdose and specimens were fixed for 2-4 days in 10% neutral buffered formalin at 4°C. Specimens were then rinsed in PBS, incubated overnight in 30% sucrose, incubated overnight in 50/50 30% sucrose and Tissue-Tek O.C.T. compound (OCT), and finally incubated overnight in OCT prior to freezing at −80°C. Cryosectioning was performed to produce 20 μm thick sections. Sections were post-fixed in 10% neutral buffered formalin and rinsed in PBS prior to further processing. Slides for fluorescent imaging were mounted with coverslips using Fluoromount G containing DAPI. Oil Red O staining was performed as previously described (38, 40). T cells were detected in sections from *TgBAC(lck:EGFP)^vcc4^* zebrafish by anti-GFP staining to enhance visible fluorescent green signal (primary antibody: ab13970, Abcam; secondary antibody: ab150173, Abcam), stained slides were then mounted with coverslips using Fluoromount G containing DAPI. All imaging was carried out on a Leica DM6000B microscope.

### Statistics

All statistical testing was carried out using Graphpad Prism. Each data point indicates a single animal unless otherwise stated.

## Acknowledgements

We thank the Centenary imaging facility core and Sydney Cytometry staff Drs Kristina Jahn, Angela Kurz, and David Liu, for their assistance.

---

**Supplemental Figure 1.**
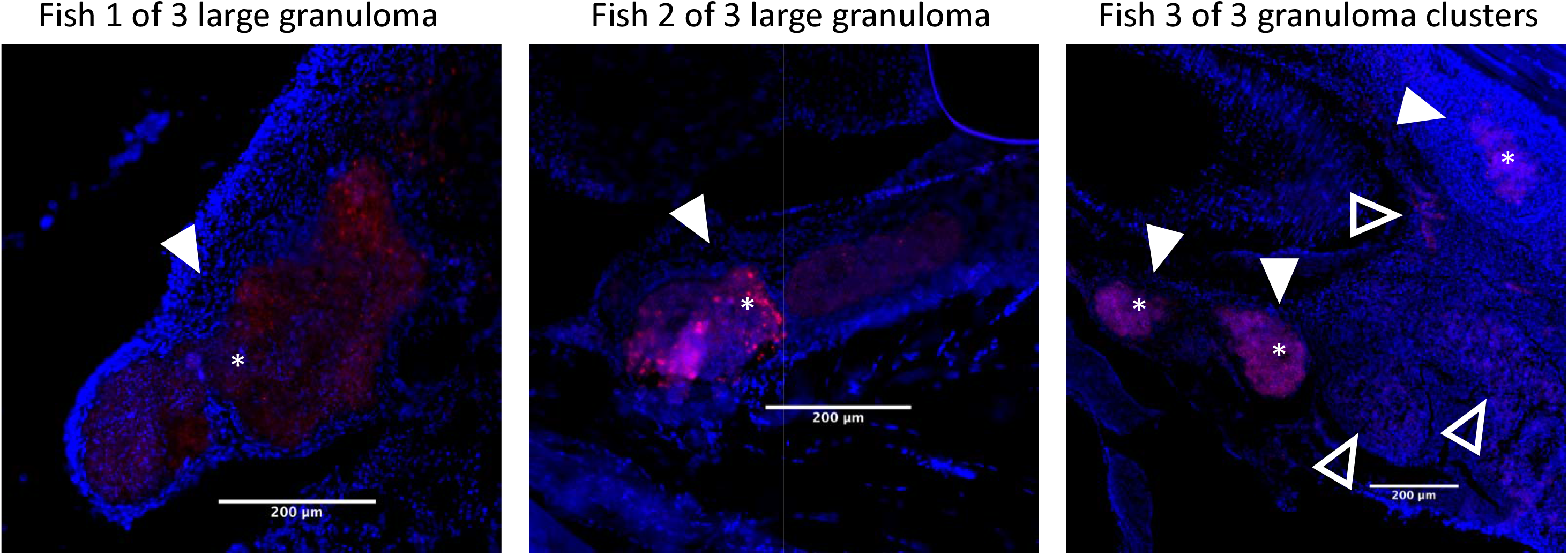
Representative images of R *M. abscessus*-tdTomato lesions in DAPI-stained cryosections from three 70 dpi adult zebrafish. Filled arrowheads indicate organised granulomas, * indicate necrotic cores, empty arrowheads indicate loose *M. abscessus* lesions. Scale bars indicate 200 μm.

**Supplemental Figure 2.**
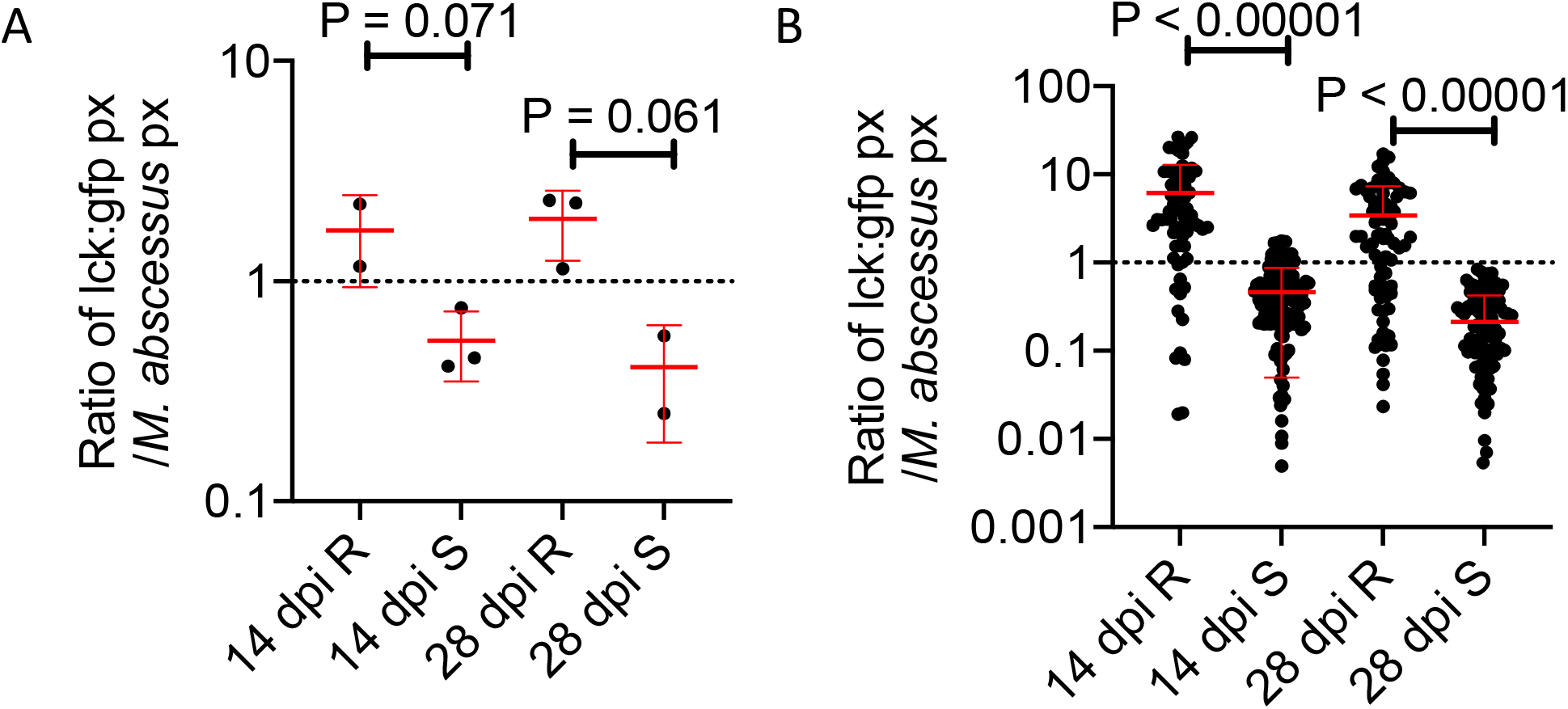
A. Quantification of T cell GFP pixels as a function of *M. abscessus*-tdTomato fluorescence in *TgBAC(lck:EGFP)^vcc4^* adult zebrafish. Each point represents the average of a single animal. Total n per group: 14 dpi R=2 S=3, 28 dpi R=3, S=2. B. Quantification of T cell GFP pixels as a function of *M. abscessus*-tdTomato fluorescence in *TgBAC*(*lck*: *EGFP*)^*vcc4*^ adult zebrafish. Each point represents a single lesion from the animals in Panel A. Total n per group: 14 dpi R=74 S=137, 28 dpi R=91, S=117.

**Supplemental Figure 3.**
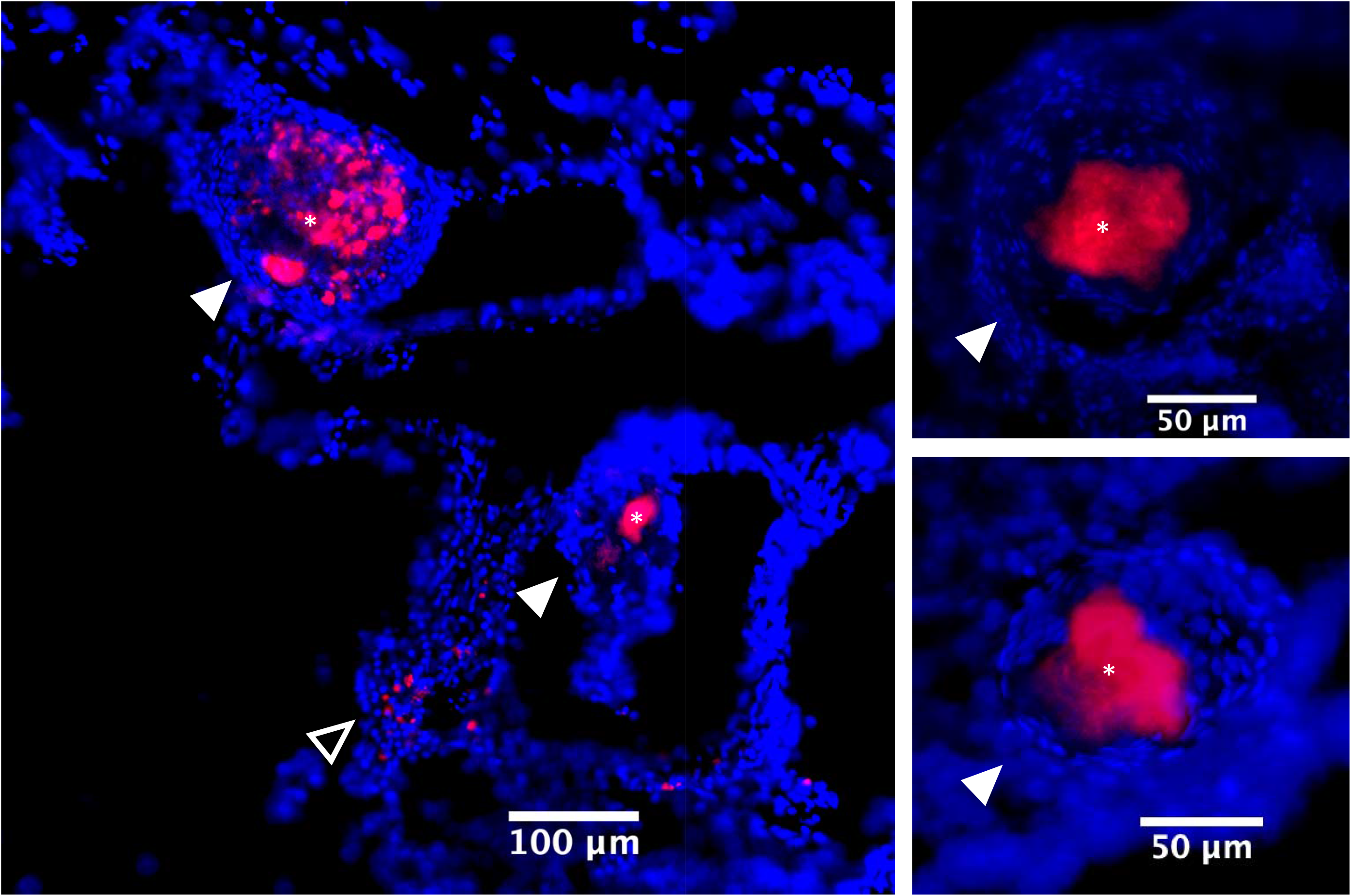
Representative images of R *M. abscessus*-tdTomato lesions in DAPI-stained cryosections from a 56 dpi *lck^-/-sa410^* fish. Filled arrowheads indicate organised granulomas, * indicate necrotic cores, empty arrowheads indicate loose *M. abscessus* lesions.

## References

1. M. D. Johansen, J. L. Herrmann, L. Kremer, Non-tuberculous mycobacteria and the rise of Mycobacterium abscessus. Nat Rev Microbiol 10.1038/s41579-020-0331-1 (2020).

2. H. Medjahed, J. L. Gaillard, J. M. Reyrat, Mycobacterium abscessus: a new player in the mycobacterial field. Trends Microbiol 18, 117–123 (2010).

3. J. F. Tomashefski, Jr., R. C. Stern, C. A. Demko, C. F. Doershuk, Nontuberculous mycobacteria in cystic fibrosis. An autopsy study. Am J Respir Crit Care Med 154, 523–528 (1996).

4. R. Nessar, E. Cambau, J. M. Reyrat, A. Murray, B. Gicquel, Mycobacterium abscessus: a new antibiotic nightmare. The Journal of antimicrobial chemotherapy 67, 810–818 (2012).

5. B. E. Ferro et al., Failure of the Amikacin, Cefoxitin, and Clarithromycin Combination Regimen for Treating Pulmonary Mycobacterium abscessus Infection. Antimicrob Agents Chemother 60, 6374–6376 (2016).

6. S. T. Howard et al., Spontaneous reversion of Mycobacterium abscessus from a smooth to a rough morphotype is associated with reduced expression of glycopeptidolipid and reacquisition of an invasive phenotype. Microbiology 152, 1581–1590 (2006).

7. E. Catherinot et al., Acute respiratory failure involving an R variant of Mycobacterium abscessus. J Clin Microbiol 47, 271–274 (2009).

8. C. R. Esther, Jr., D. A. Esserman, P. Gilligan, A. Kerr, P. G. Noone, Chronic Mycobacterium abscessus infection and lung function decline in cystic fibrosis. J Cyst Fibros 9, 117–123 (2010).

9. B. E. Jonsson et al., Molecular epidemiology of Mycobacterium abscessus, with focus on cystic fibrosis. J Clin Microbiol 45, 1497–1504 (2007).

10. A. Bernut et al., Mycobacterium abscessus cording prevents phagocytosis and promotes abscess formation. Proc Natl Acad Sci U S A 111, E943–952 (2014).

11. H. Medjahed, J. M. Reyrat, Construction of Mycobacterium abscessus defined glycopeptidolipid mutants: comparison of genetic tools. Appl Environ Microbiol 75, 1331–1338 (2009).

12. A. V. Gutierrez, A. Viljoen, E. Ghigo, J. L. Herrmann, L. Kremer, Glycopeptidolipids, a Double-Edged Sword of the Mycobacterium abscessus Complex. Front Microbiol 9, 1145 (2018).

13. A. Bernut, J. L. Herrmann, D. Ordway, L. Kremer, The Diverse Cellular and Animal Models to Decipher the Physiopathological Traits of Mycobacterium abscessus Infection. Front Cell Infect Microbiol 7, 100 (2017).

14. A. Bernut et al., In vivo assessment of drug efficacy against Mycobacterium abscessus using the embryonic zebrafish test system. Antimicrob Agents Chemother 58, 4054–4063 (2014).

15. A. Obregon-Henao et al., Susceptibility of Mycobacterium abscessus to antimycobacterial drugs in preclinical models. Antimicrob Agents Chemother 59, 6904–6912 (2015).

16. M. Rottman et al., Importance of T cells, gamma interferon, and tumor necrosis factor in immune control of the rapid grower Mycobacterium abscessus in C57BL/6 mice. Infect Immun 75, 5898–5907 (2007).

17. C. T. Oh, C. Moon, M. S. Jeong, S. H. Kwon, J. Jang, Drosophila melanogaster model for Mycobacterium abscessus infection. Microbes Infect 15, 788–795 (2013).

18. M. Meir, T. Grosfeld, D. Barkan, Establishment and Validation of Galleria mellonella as a Novel Model Organism To Study Mycobacterium abscessus Infection, Pathogenesis, and Treatment. Antimicrob Agents Chemother 62 (2018).

19. A. Bernut et al., Mycobacterium abscessus-Induced Granuloma Formation Is Strictly Dependent on TNF Signaling and Neutrophil Trafficking. PLoS Pathog 12, e1005986 (2016).

20. V. Dubois et al., MmpL8MAB controls Mycobacterium abscessus virulence and production of a previously unknown glycolipid family. Proc Natl Acad Sci U S A 115, E10147–E10156 (2018).

21. I. Halloum et al., Deletion of a dehydratase important for intracellular growth and cording renders rough Mycobacterium abscessus avirulent. Proc Natl Acad Sci U S A 113, E4228–4237 (2016).

22. A. Bernut et al., CFTR Protects against Mycobacterium abscessus Infection by Fine-Tuning Host Oxidative Defenses. Cell reports 26, 1828–1840 e1824 (2019).

23. C. A. Madigan, J. Cameron, L. Ramakrishnan, A Zebrafish Model of Mycobacterium leprae Granulomatous Infection. J Infect Dis 216, 776–779 (2017).

24. S. H. Oehlers et al., Interception of host angiogenic signalling limits mycobacterial growth. Nature 517, 612–615 (2015).

25. L. E. Swaim et al., Mycobacterium marinum infection of adult zebrafish causes caseating granulomatous tuberculosis and is moderated by adaptive immunity. Infect Immun 74, 6108–6117 (2006).

26. M. Parikka et al., Mycobacterium marinum Causes a Latent Infection that Can Be Reactivated by Gamma Irradiation in Adult Zebrafish. PLoS Pathog 8, e1002944 (2012).

27. L. Marjoram et al., Epigenetic control of intestinal barrier function and inflammation in zebrafish. Proc Natl Acad Sci U S A 112, 2770–2775 (2015).

28. K. Sugimoto, S. P. Hui, D. Z. Sheng, M. Nakayama, K. Kikuchi, Zebrafish FOXP3 is required for the maintenance of immune tolerance. Dev Comp Immunol 73, 156–162 (2017).

29. F. M. Collins, Mycobacterial disease, immunosuppression, and acquired immunodeficiency syndrome. Clin Microbiol Rev 2, 360–377 (1989).

30. A.-L. Roux et al., The distinct fate of smooth and rough Mycobacterium abscessus variants inside macrophages. Open biology 6 (2016).

31. A. Bernut et al., Mycobacterium abscessus cording prevents phagocytosis and promotes abscess formation. Proceedings of the National Academy of Sciences 111, 943–952 (2014).

32. T. Mogues, M. E. Goodrich, L. Ryan, R. LaCourse, R. J. North, The relative importance of T cell subsets in immunity and immunopathology of airborne Mycobacterium tuberculosis infection in mice. J Exp Med 193, 271–280 (2001).

33. J. D. Yang et al., Mycobacterium tuberculosis-specific CD4+ and CD8+ T cells differ in their capacity to recognize infected macrophages. PLoS Pathog 14, e1007060 (2018).

34. V. Le Moigne et al., Efficacy of Bedaquiline, Alone or in Combination with Imipenem, against Mycobacterium abscessus in C3HeB/FeJ Mice. Antimicrob Agents Chemother 64 (2020).

35. D. X. Ji et al., Type I interferon-driven susceptibility to Mycobacterium tuberculosis is mediated by IL-1Ra. Nat Microbiol 4, 2128–2135 (2019).

36. I. K. Park et al., Clonal Diversification and Changes in Lipid Traits and Colony Morphology in Mycobacterium abscessus Clinical Isolates. J Clin Microbiol 53, 3438–3447 (2015).

37. A. Pawlik et al., Identification and characterization of the genetic changes responsible for the characteristic smooth-to-rough morphotype alterations of clinically persistent Mycobacterium abscessus. Mol Microbiol 90, 612–629 (2013).

38. T. Cheng, J. Y. Kam, M. D. Johansen, S. H. Oehlers, High content analysis of granuloma histology and neutrophilic inflammation in adult zebrafish infected with Mycobacterium marinum. Micron 129, 102782 (2020).

39. A. Bernut et al., Deciphering and Imaging Pathogenesis and Cording of Mycobacterium abscessus in Zebrafish Embryos. J Vis Exp 10.3791/53130 (2015).

40. M. D. Johansen et al., Mycobacterium marinum infection drives foam cell differentiation in zebrafish infection models. Dev Comp Immunol 88, 169–172 (2018).

